# RNAGAN: Train One and Get Four, Multipurpose Human RNA-Seq Analysis Tool with Enhanced Interpretability and Small Data Size Capability

**DOI:** 10.64898/2026.03.17.712527

**Authors:** Zhaozheng Hou, Victor Ho-Fun Lee, Dora Lai-Wan Kwong, Xinyuan Guan, Zhonghua Liu, Wei Dai

**Affiliations:** Department of Clinical Oncology, University of Hong Kong, Hong Kong (SAR), PR China; School of Biomedical Sciences, University of Hong Kong, Hong Kong (SAR), PR China; Department of Biostatistics, Columbia University, New York, USA; Department of Clinical Oncology, Shenzhen Key Laboratory for cancer metastasis and personalized therapy, The University of Hong Kong-Shenzhen Hospital, Shenzhen, PR China

## Abstract

The advent of artificial intelligence (AI) has brought revolutionary tools for biomedical transcriptomic (RNA-level) research. However, there are persistent constraints including limited interpretations with biomedical concepts such as functional pathways, small sample sizes and substantial time and computing power requirements for AI training.

To overcome these limitations, we developed RNAGAN (https://github.com/ZhaozhengHou-HKU/RNAGAN-1.0.git), an AI tool with a generative adversarial network (GAN) structure with the objective of enhancing transcriptomic analysis. The network was established based on public human datasets comprising 4.6 million single cells from multiple organs and 5,900 sequenced samples of various cancer types with normal references. A specialized pathway neural layer was embedded to extract activities of predefined pathways from the Human Molecular Signatures Database (MSigDB), or newly learned pathways from single-cell data.

The structure of RNAGAN (generator and discriminator) enables four applications after one shared training procedure: 1. single-cell and bulk-level patient stratification or differential diagnosis; 2. analysis of the gene and pathway markers in a selected disease; 3. pseudo data generation when sample size is limited for downstream analysis; 4. vectorization with gene and pathway-level features learned from multiple data sets. RNGAN contributes to the efficient utilization of limited data for transcriptomic studies.

## Introduction

Single-cell and bulk RNA sequencing have revolutionized our understanding of cellular heterogeneity, tissue complexity, and disease mechanisms [1–4]. However, the high dimensionality and inherent sparsity of transcriptomic data pose significant challenges for robust stratification, mechanistic interpretation, and meaningful comparison among samples. Moreover, current analysis pipelines involve batch-level normalization followed by differential expression (DE) analysis, which require a relatively large sample size, limiting their applicability in scenarios with a paucity of reference samples, such as rare diseases, newly recognized phenotypes, and policy constraints [5].

Meanwhile, advancements in artificial intelligence (AI) approaches have significantly extended the capabilities of RNA sequencing and multiple omics. Recent AI tools mainly focused on single-cell-level analysis, such as: scDesign3 [6], scPRAM [7], scRegulate [8], CelLink [9], CellScope [10], while there are less tools available for bulk sequencing, such as SimBu [11] which can generate high quality pseudo bulk RNA sequencing results from high-throughput single-cell transcription data. Furthermore, there are tools for multi-omics data, which has the protentional of reveal more comprehensive mechanisms [12–17], yet there are also concerns about practical constrains of data availability such as the and affordability in clinical applications [18] and ownership of the data [19], which could pose significant constraints.

A serious concern for current AI applications is their accuracy and interpretability [20, 21]. Multiple benchmark studies show that many of the AI models available are not yet accurate enough for practical clinical applications and may even be inferior to classical approaches (such as linear models or classical statistical analysis) in some cases [22–24]. Regarding to one of the most representative global medical projects in 2025, virtual cell model challenge [25], two major appeals of future work have been identified as: a more thorough analysis and verification with wet-lab experiments, and the embedding of current medical knowledge in the establishment of AI models [26, 27]. This issue, once again, connects to the medical challenge previously mentioned, which pertains to the limited amount of corresponding experimental data in the wet laboratory because of numerous practical difficulties.

Regarding generative AI tools, an additional concern is the risk of data memorization, whereby a model may inadvertently retain information from specific real samples and subsequently generate outputs that are nearly identical “pseudo samples.” In clinical applications, this could constitute a grave infringement of privacy protections [28]. This phenomenon can also result in an overestimation of the model’s performance [20]. This challenge has yet to be comprehensively addressed, which requires meticulous consideration in the development of AI models [29].

To address these challenges, we propose a novel generative adversarial network (GAN) framework designed specifically for transcriptomic data (single-cell or bulk RNA sequencing). Our approach serves four key purposes. First, it enables cell or sample-level stratification and diagnostic inference (such as cancer sample vs negative reference, or groups of different phenotypes or clinical outcomes) using as few as 20–30 positive reference samples. Second, it facilitates gene- and pathway-level interpretation of stratification results or difference between sample groups, identifying key marker genes and potential regulatory mechanisms underlying observed patterns. Third, the model can generate high-quality pseudo samples, enhancing data richness and improving sample representativeness for downstream analyzes. Fourth, generate the vectors corresponding to a given sample population, which can be used to search for similar or related populations for downstream analysis such as differential expression analysis. The model is available at https://github.com/ZhaozhengHou-HKU/RNAGAN-1.0.git.

This study explored multiple new strategies to achieve multiple functions with one AI model and one training procedure, structurally embed widely recognized medical knowledge (particularly about pathways) into the model, enhance performance with limited sample size, and prevent the memorization problem. These designs and concepts could be adopted in future AI networks for medical applications.

## Materials and methods

### Data collection and preprocessing

In this study, we used RNA sequencing data from human samples [21]. We collected 14 single-cell RNA-seq datasets of different diseases and tissues (4.6M cells in total) via CZ CELLxGENE Discover [22], and 8 bulk RNA-seq real human sample datasets of different diseases and tissues (5.9 thousand sequenced samples in total) from the Cancer Genome Atlas (TCGA) database or the Gene Expression Omnibus (GEO) database [23, 24]. All datasets used are publicly available (details are summarized in **Table S1** and **S2**).

70% of the samples were used for network training, and 30% were used for validating the performance of the network. For single-cell dataset, this training-validation division applied to each cell type group from each patient; for bulk-sequencing dataset, this applied to every sample group for each detailed disease type from each dataset. In training and validation, the references for RNAGAN were generated by random sampling (with replacement) the required number of samples from the corresponding sample group. The groups with less than five members were not adopted.

The gene expression level was normalized in the form of Fragments Per Kilobase per Million mapped fragments (FPKM) [25], which considers the total length of exons of genes and the total sequencing depth. This method has been chosen because:

1. it improves the comparability between different genes, which is beneficial for learning pathways and carrying out pathway enrichment analysis;
2. it makes the expression level for single-cell data and bulk-sequencing data more comparable, which leads to a smoother task transfer for RNAGAN;
3. it does not correct the cohort or population level features, which allows the RNAGAN to learn natural variances patterns among cohorts, thus exhibiting more stable performance across different datasets (particularly new datasets in protentional applications).

We noticed that different datasets cover different genes. To achieve a balance between generality and model training performance, we only selected coding genes that are captured by at least 50% of single-cell datasets (18,583 genes, about 94.0% of all appeared coding genes), and the expression level of missing genes were set to 0.

### Pathway Information

We investigated the performance of networks processing the pathway-level features, and one of the approaches is using predefined pathways. The Human Molecular Signatures Database (MSigDB, v2023.1.Hs, Mar 2023) was adopted as the reference [26, 27, 30], and the following four signature sets were selected: Hallmark gene sets (50 pathways), KEGG legacy sets (186 pathways) [28], Gene Ontology (10,454 pathways): biological process (BP), cellular component (CC), and molecular function (MF), and Reactome gene sets (1,736 pathways).

To filter out duplicate or highly similar pathways, pairwise Jaccard distances between these pathways were calculated, and for pairs with distance less than 0.5 (i.e., more than 50% of pathway elements are common) only the one with more gene members would remain. With this filtering standard, there were 8,599 pathways remained to be embedded in the GAN network as the predefined pathways (the corresponding list is available in Table S5).

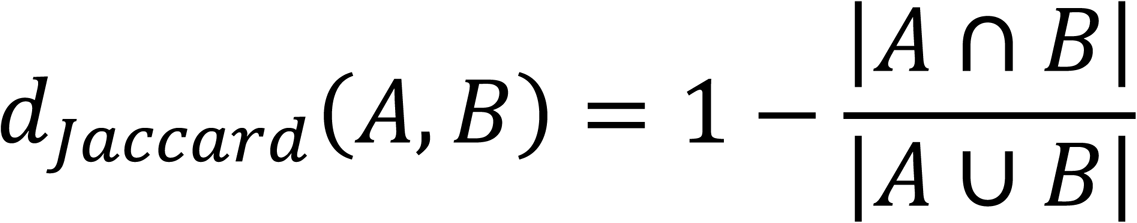

### Network design

We used MATLAB 2024b Deep Learning Toolbox to establish the model and implement some customized layers (such as data-split layer for splitting the sample to diagnose and the reference data, which can also be achieved with an equivalent standard convolution layer). We carried out hyperparameter optimization with grid search of network structure/parameter combinations: pathway feature extraction = NP-No Pathway, PP-Predefined Pathways, or LP-Learnable Pathways; number of references = 10, 20, or 30. Number of learnable and layers for sub-networks are given in **Table 1**.

**Table 1.** Number of learnable and layers for sub-networks.

| Pathway features | Generator network<br># of learnable and layers | Discriminator network<br># of learnable and layers |
| --- | --- | --- |
| NP-No Pathway | 19.1M (33 layers) | 14.3M (21 layers) |
| PP-Predefined Pathways | 183.3M (35 layers) | 180.7M (24 layers) |
| LP-Learnable Pathways | 183.3M (35 layers) | 180.7M (24 layers) |

As mentioned in the introduction, the GAN structure involves two sub-networks: the Generator network and the Discriminator network, and the corresponding network structures are shown in **Figure 1**. The detailed design and explanation of the network is given as follows, using the Generator of PP20 (predefined pathways, using 20 references for data generation) as an example:

**Figure 1.**
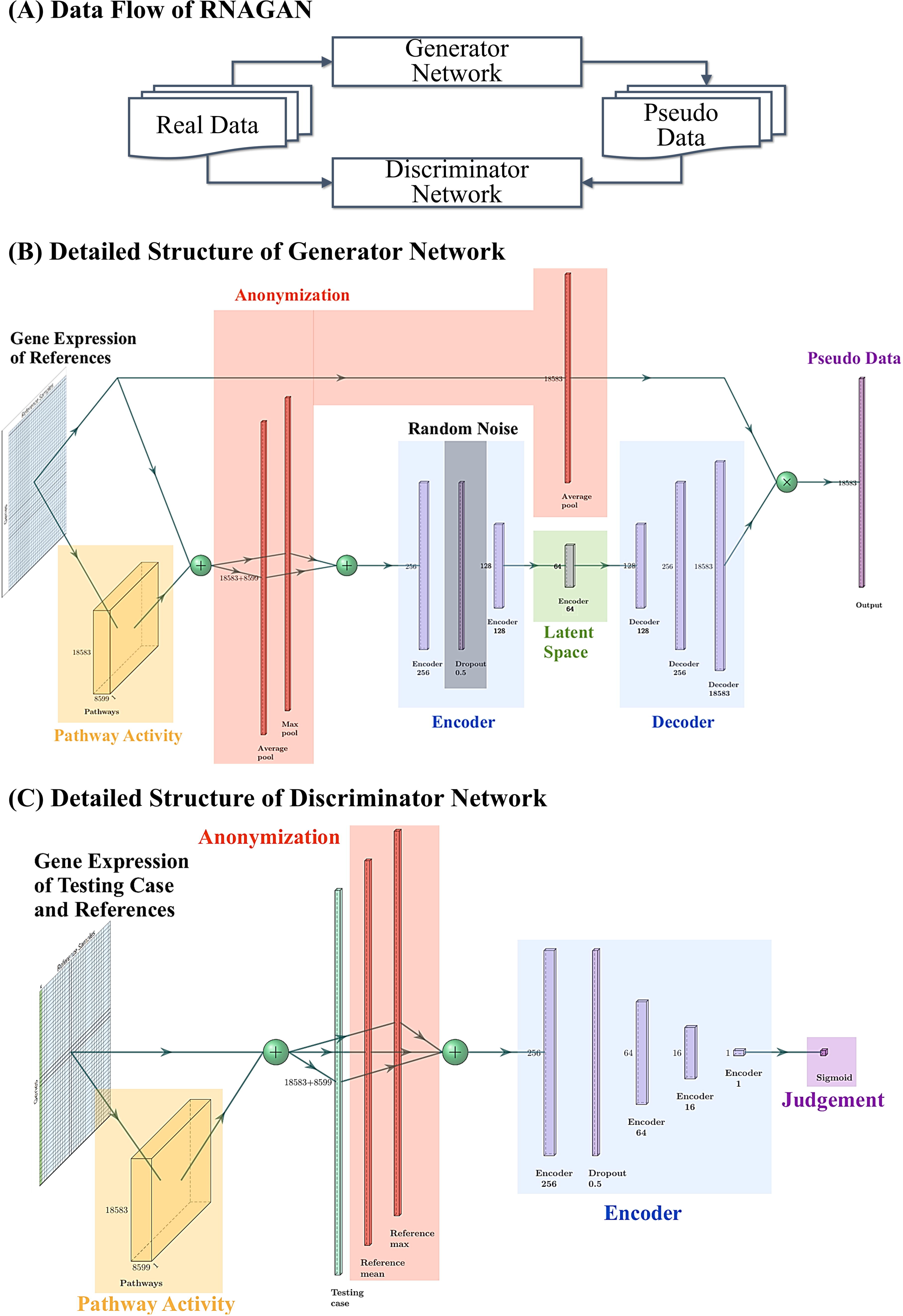
RNAGAN structure. (**A**) The RNAGAN is established with two sub-networks: the generator network and the discriminator network. The generator’s task is to generate pseudo data that is similar to the given real data, and the discriminator’s task is to differentiate pseudo data (or real data from different sample groups) from real data. (**B**) structure of the generator labeled with functions. (**C**) structure of the generator labeled with functions.

### Major network layers

i. **Input layer:** The PP20 generator network uses the imageInputLayer to load the input, with a size of 18,583 by 20 by 1, with each “pixel”" being a single-precision floating-point number corresponding to the FPKM for each gene in each reference sample. Although the data are not a real or meaningful image, the imageInputLayer is adopted to simplify the data preparation and follow-up process; it also demonstrates a simple and effective approach to establishing specialized networks with a general-purpose machine learning toolbox;
ii. **Pathway layer:** The input data is then used to calculate the pathway activity, which is achieved by a specialized convolution layer. The layer has 8,599 channels (corresponding to the number of pathways) with a filter size of 18,583 by 1, so that the weight values of the filter correspond to the weight of each gene in each pathway (0 and 1 for the networks with predefined pathways);
iii. **Anonymization layers:** The results of gene expression and pathway activity are then concatenated into one table, and the mean and maximum values among all samples are calculated by the two corresponding pooling layers. These two layers are typical machine learning layers (i.e., well-optimized for data processing efficiency) and can reflect the average expression/activity level and variance within the references. More importantly, these layers structurally ensure that the network cannot forward exact expression values of any individual sample as its output, which avoids low-quality copy-pasting pseudo data and leakage of the exact gene expression of any single sample.
iv. **Reasoning U-net:** A U-net-like structure is adopted to analyze the gene and pathway features to identify the deeper features/patterns of the referencing population, and then this information is used to generate pseudo gene-expression data. The U-net reduced the channels to 64 with multiple layers and then decoded them back to 18583 (the number of genes). A dropout layer is added to introduce noise to the pseudo data and improve the robustness of the final model.
v. **Multiplication Layer:** Multiply the U-net output by the average gene expression of the references, and the final product values will serve as the network output. This layer addresses two considerations: it structurally avoids “spurious values” of unavailable genes (i.e., if all references have a zero reading for a gene or if that gene is not available in the dataset, the pseudo data should also yield 0 for that gene); it reduces the difficulties faced by U-net, as the gene expression level of the pseudo data should be comparable to the average expression level on scale. With this design, the training/learning of the network can focus more on the pattern of natural variance within populations than on the reading sensitivity of a certain gene with a specific sequencing assay.

### RNAGAN architecture variants

i. **Different Number of References:** The widths of the input data table correspond to the number of references. Changes in the amount of data to be used in applications does not change the number of network parameters.
ii. **Discriminator Network:** It uses the input with a width of 1+n (the first column is the sample to discriminate). It shares most of the structures with the generator but only has the encoder side of the U-net followed with a sigmoid output layer. The output ranges from 0 to 1, and a score close to 0 indicates that the testing sample has some unique features and is unlikely to be of the same sample type as the given reference.
iii. **NP and LP networks:** The NP (No Pathway) version does not have the pathway layer and directly forwards gene-level information to the U-net; LP (Learnable Pathways) version uses the same number of pathways/channels (8,599) and initializes the pathway layer with randomized weight. The LP allows the layer to learn during training with single-cell data, and then the weight is fixed when network training progresses to bulk-sequencing data because Bulk-sequencing data can “pollute” the layer with niche/microenvironment-level features (such as cell type proportion-related features rather than biomedical functions) that would make the learned features difficult to guide further biomedical studies (such as functional studies) as identified pathways.

### Network training

We trained the GAN networks with hyperparameter grid search (nine combinations in total), with the negative log-likelihood loss. The corresponding loss function is as follows:

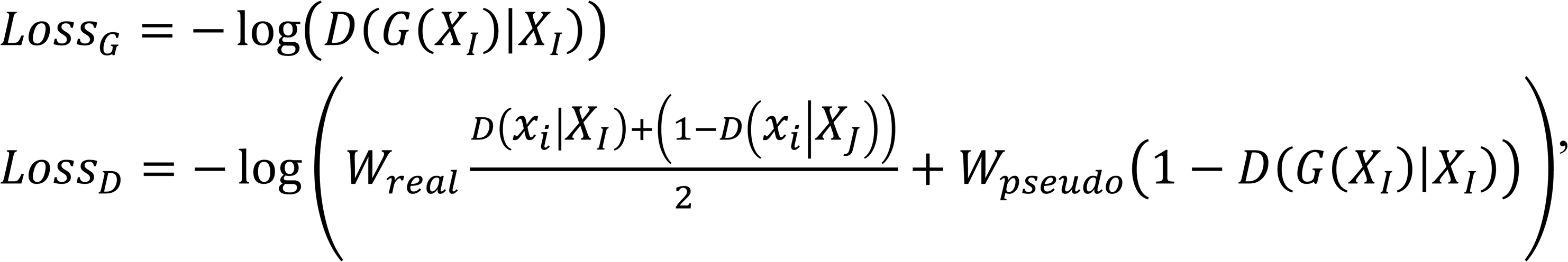

where *X*_*I*_ and *X*_*J*_ are two different real sample groups, *x*_*i*_ is an element from group *X*_*I*_, *G*(*X*_*I*_) is generated pseudo results using *X*_*I*_ members as reference, *D*(*x*_*i*_|*X*_*I*_) is discrimination results of whether *x*_*i*_ is an element from group *X*_*I*_, *W*_*real*_ and *W_pseudo_* are the evaluation weight of discriminator’s performance on real and pseudo data. Since the discriminator may be used by its own with real data in clinical diagnosis tasks, and identifying pseudo data is not really a clinically relevant application, we set the weight as *W*_*real*_: *W_pseudo_* = 10: 1.

As mentioned in the introduction, the training process has been divided into three steps and the corresponding settings are given in **Table 2**. The learning rates (the “step length” of the network variable change during training) were set as given so that: the discriminators could quickly establish a valid understanding of the data in step 1 (as pseudo data from early stage generators could be unhelpful and unhelpful for discriminator training) [31], then generators caught up with the discriminators to achieve high-quality pseudo data with a higher learning rate (to make sure generators could “win” the competition against discriminators, otherwise generators could have model collapsing problem being overpowered by discriminators [29]), then bulk-sequencing data was given to the networks with a lower learning rate as a transfer learning stage. It is a Model-Based Transfer Learning (MBTL), also a cross-domain transfer learning procedure. The learning rate was restarted at each step as suggested by Li & Kang [32]. The numbers of epochs (times for each sample being used in network training) were large enough to observe that the performance on validation dataset stops improving in validation for most of network structure/parameter combinations.

**Table 2.**
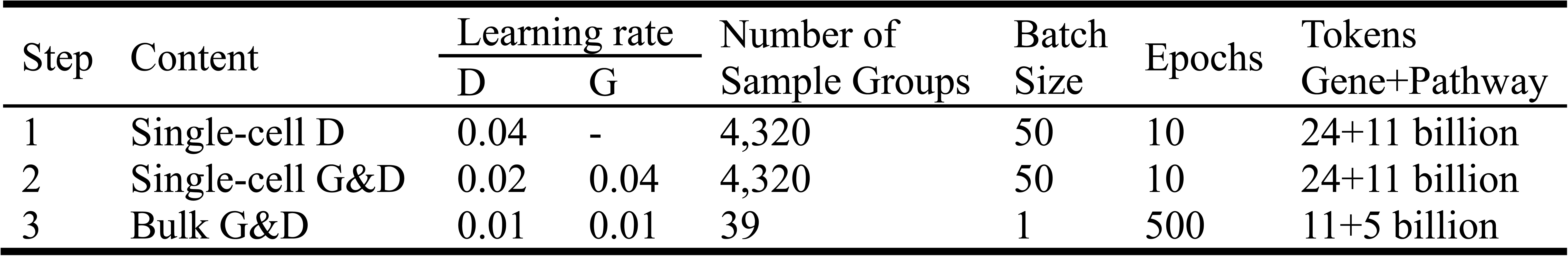
Training settings.

### Vectorization and visualization of bulk-seq sample groups

The generator of RNAGAN can provide the latent space vector corresponding to a given set of samples. The network reduce embeds the expression pattern of 18,583 genes of the sample set into a 64-dimensional vector, which can be further reduced to 2 or 3 dimensions with PCA, UMAP, or other similar tools for visualization. In this example, we used PCA for this purpose and compared with the dimension reduction results directly from the averaged gene expressions, with common vectorization methods including PCA, UMAP and T-SNE [33].

## Results

The RNAGAN structure can be split into two sub-networks: Generator network and Discriminator network, which can support different analysis tasks. The detailed deliverables from the networks, and the features available for interpretation are given in **Figure 2**. As shown in the figure, gradient-weighted activation mapping (Grad-CAM) [34, 35], occlusion sensitivity [36, 37], and gradient-based attribution [38] were adopted to extract the features identified from the discriminator networks.

**Figure 2.**
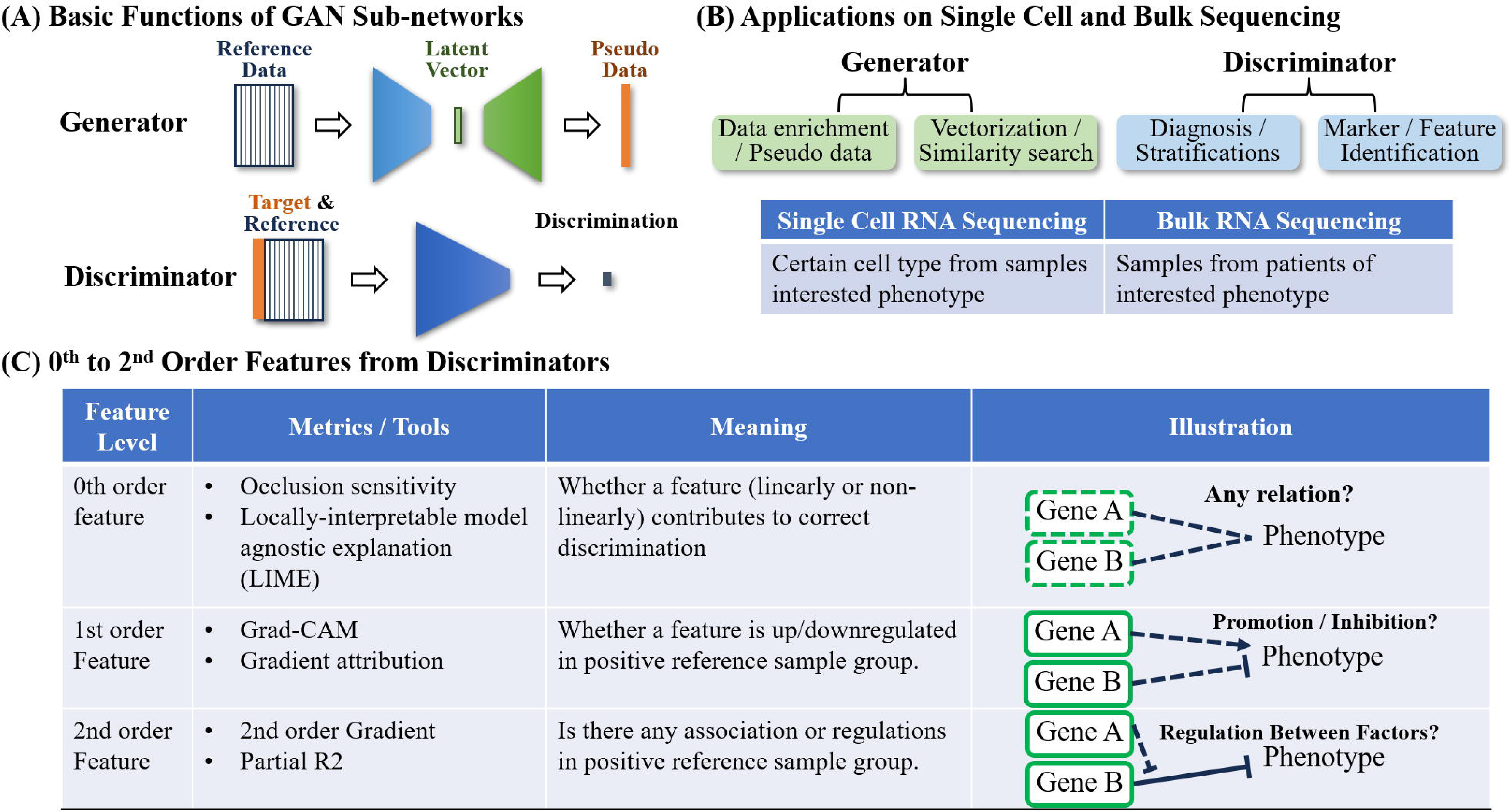
Results that RNAGAN networks can provide. (**A**) Basic Functions of GAN Sub-networks. (**B**) Detailed results of the generator and discriminator with single-cell and bulk RNA sequencing data. (**C**) Features that can be extracted from the discriminator for the interpretation of the output of these networks.

### Function 1: Patient Stratification and Differential Diagnosis

The Discriminator can be used for general purpose identification of cell types or diseases diagnosis with given reference samples. With prepared positive and negative referencing samples (such as breast cancer vs. normal samples, or classical glioblastoma vs. mesenchymal glioblastoma) and one sample to diagnose, discriminator can compute the discrimination score to quantitively analyze whether the sample is more similar to the positive references or negative ones.

For each trained network, a series of evaluation metrics have been adopted to quantify its performance: area under the curve (AUC), accuracy, and F1-score. The evaluations are based on the validation data in all sample groups (4,320 cell type-patient groups for single-cell applications, and 39 disease subtype groups for bulk-level applications), 10 random sampling trials for each group. The average score is shown in **Figure 3**.

**Figure 3.**
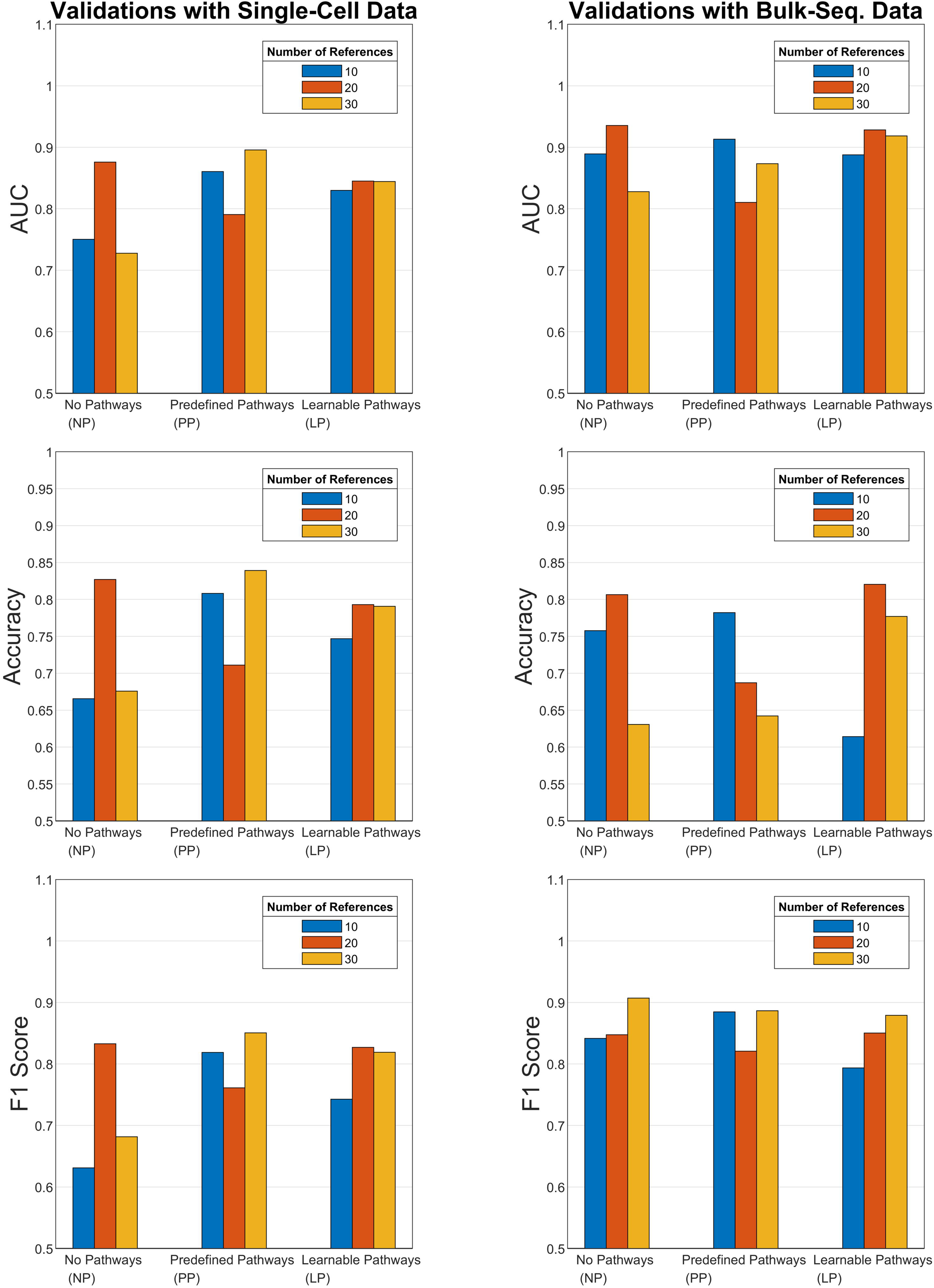
Weighted average scores for all discriminator networks.

As shown in **Figure 3**, Performance in validation data achieved averaged area under the curve (AUC) over 70% in cell type identification with single-cell RNA sequencing, and over 80% averaged AUC in diseases diagnosis with bulk-sequencing. One of the reasons to explain the difference of performance between single-cell and bulk RNA sequencing is that there are many more cell subtypes based on single-cell data, which make diagnosis a more difficult task.

For discriminators with learnable pathways (LP), the network with 20 or 30 references outperforms the 10-reference version in both single-cell and bulk applications, which suggests that informativeness of pathways learned during the training depends on the number of references available, and 10 reference samples may not be sufficient in general. For the application on bulk RNA sequencing, validation AUC of LPs with 20 or 30 references achieved above 90%.

To facilitate researchers working on specific cancer types, such as Breast Cancer, Glioblastoma, and Malignant Melanoma, we applied our RNAGAN model to 13 distinct cancers with more than 30 bulk-RNA sequencing samples. The diagnosis accuracy evaluations (**Table S3**) and identified gene/pathway-level markers are detailed in **Table S4**.

In the accuracy evaluations in terms of AUC (**Table S3**), we noticed that the performance is significantly affected by the number of available samples the cancer types with more than 20 validation references generally lead to AUC higher than 80% in the models of NP20 and PP20. On the other hand, for some cancer types including classic glioblastoma multiforme (GBM) and acute myeloid leukemia (AML), some models achieved AUC scores over 75% while the number of available references is less than 20. It may be because these conditions have significant biomedical differences compared to the negative samples, thereby allowing the model to effectively leverage these distinctions for diagnostic purposes.

### Function 2: Explanation designed for medical professionals

Feature explanation was extracted from the discriminators for the purpose of providing explicit reasonings of the discrimination results and providing hints to medical researchers about the gene/ path-level features that are related to the difference between samples. With corresponding metrics and tools as shown in **Figure 2C**, it can be quantitatively analyzed that: whether the information of a certain gene/pathway contributes to a correct discrimination (0^th^ order feature), is the gene/pathway is more enriched in positive or negative reference samples (1^st^ order feature), and if one of the gene/pathway has associations with another gene or dominates the diagnosis against the contribution of another gene/pathway (2^nd^ order feature).

RNA sequencing results (also the network input) were normalized into FPKM values, which consider the total length of exons of genes and the total sequencing depth. Batch-level correction was not included because RNAGAN was designed to be capable with small data size (n≈30), and multiple recent studies have revealed the considerable risk of mis-correction and true signal loss with limited samples [39–41]. However, samples from the same platform and pipeline as a reference are strongly recommended. Supported sample group pairing strategies are given in **Table 3**.

**Table 3.** Supported bulk-RNA sequencing sample group pairs for feature extraction.

| Positive vs. Negative Reference | Example |
| --- | --- |
| Certain subgroup of a disease vs. other subgroups of the same disease | Classical Glioblastoma vs. Proneural and/or Mesenchymal Glioblastoma;<br>Breast Carcinoma with metastasis vs. without metastasis |
| Samples of certain disease vs. comparable normal samples | Infiltrating Lobular Breast Carcinoma vs. adjacent normal samples |
| Samples of certain disease vs. other diseases from the same study dataset | Breast Carcinoma vs. other diseases from the same dataset |

We present the gene and path-level features that have been extracted from function 1 applications in the supplementary **Table S4**. The features were selected from the NP and PP models with the highest Area Under the Curve (AUC). LP for Classical GBM was employed rather than PP because LP models perform significantly better than PP in that case. The standard for feature extraction was as follows: the median and mean of occlusion sensitivity had to be positive among the 100 positive (targeted samples as the "unknown") and 100 negative trials (negative samples as the "unknown") with a randomly selected subset of targeted samples (different from the "unknown" on for positive cases) as the reference. The average gene/ path activity of the targeted samples had to have an absolute log2 fold change larger than 1 compared to negative trials. In instances involving more than 50 filtered features, a ranking system was used to prioritize features based on the magnitude of the fold change, resulting in the selection of the top 50 features.

To interpret the self-learned pathways from the LP versions to users, following one of the most widely adopted gene set enrichment analysis (GSEA) [26], we used the Kolmogorov–Smirnov test for enrichment analysis to identify which pathways or functions get reflected by which self-learned pathway. The learned pathways were defined as a set of gene-weight pairs that have been extracted from the pathway neural layer. The genes were then sorted according to the given weight and compared to predefined pathways (i.e., gene sets) to ascertain whether related genes were significantly enriched or depleted. **Table S5** provides the predefined pathway list for PP networks, and the top 5 enriched and depleted pathways for each learned features in LP networks, so that pathway features from LP can be "translated" into known pathways or mechanisms.

It can be posited that the extracted features are meaningful within the context of the current medical understanding of the specific cancer type. For instance, we carried out a comparison made between two widely adopted and validated gene testing tools for breast cancer: Oncotype DX [42] and MammaPrint [43]. The related genes in the toolkits for diagnosing are significantly enriched in the selected top 50 marker genes and pathways (p=0.042 for Oncotype, and p=0.015 for MammaPrint), such as MMP11, GRB7, and ERBB2 from Oncotype, and WISP1, PRAME, and RAD21 from MammaPrint.

The extracted features for Infiltrating Lobular Breast Carcinoma (ILC) are shown in **Figure 4**. Using gene WISP1 and MPO as examples, they both have made significant contribution to the correct diagnosis of RNAGAN (0^th^ order feature), WISP1 is a positive marker of cancer samples, and MPO is the opposite (1^st^ order feature), and WISP1 dominates the cancer sample discrimination as the samples expressing both MPO and WISP1 are more likely to be a BCILC case (2^nd^ order feature). WISP1 (WNT1-inducible signaling pathway protein 1) is a well-known oncogene, promoting tumor growth, metastasis, and epithelial-to-mesenchymal transition (EMT) in ILC [44], which is also reported to be correlated to infiltration of immune cells including neutrophils [45]. MPO (Myeloperoxidase) is highly related to adaptive immunity, and the infiltrated MPO-positive cells help to reduce tumor size and improve overall survival [46, 47]. These findings are consistent with our analysis results for these two important marker genes in breast cancer.

**Figure 4.**
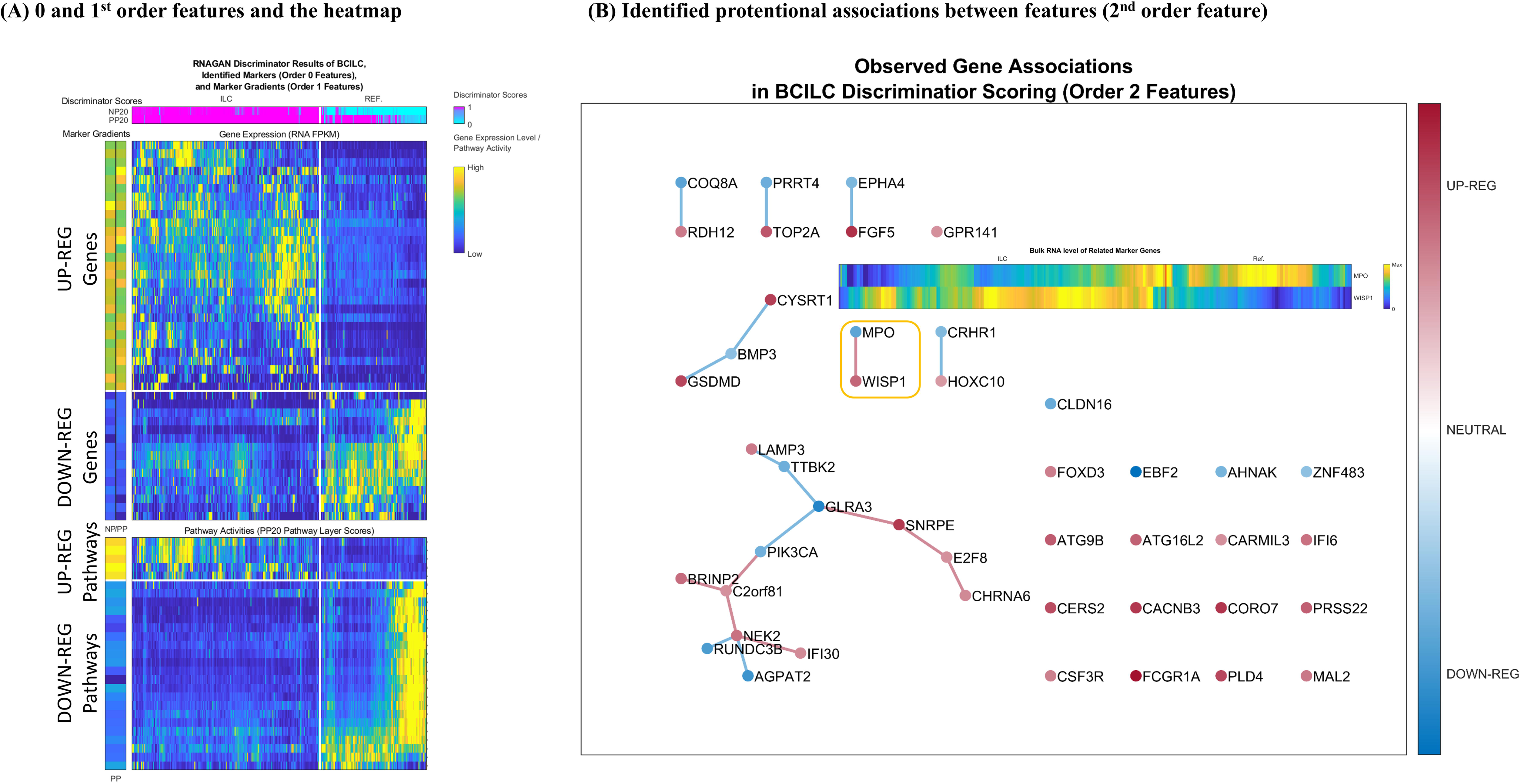
Feature extraction for Infiltrating Lobular Breast Carcinoma (ILC) from discriminators in versions NP20 and PP20. (**A**) 0 and 1st order features and the heatmap showing these genes and pathways are significantly correlated to the comparison of ILC vs. normal reference. (**B**) Identified protentional associations between gene-level features. It should be emphasized that these results are purely data-driven and require further analysis and experiments before clinical applications.

The identified features in **Figure 4** are not only significant for differential expression in raw data but also supported by the RNAGAN network as key evidence to make discrimination between cancer and normal samples. There are some other features that may be statistically significant according to the raw data but rejected by the RNAGAN, because the GAN network learned from other datasets that there could be considerable level of within-group variance in these features and thus these features may not be as stable and robust as the remaining features. The generated 2^nd^ order feature plots for breast cancer ILC and classical GBM were illustrated in **Figure S1** as examples.

### Function 3: Data synthesis/enrichment

The generator of RNAGAN is designed for generating pseudo data bases on given references in the form of FPKM. To validate the quality of synthetic bulk sequencing data, we compared the results of all generators with scDesign3 [6] and an approach based on Bayesian statistics [48] as baselines. The corresponding results are shown in **Figure 5**. The pseudo data generated from all the generators have significantly better quality in terms of passing the discrimination from the discriminator networks (P<0.001, Wilcoxon tests). In this analysis, the values for the paired discriminator against the generator were not suitable for comparison because they interacted with each other during the training procedure. The corresponding values have been replaced with the average score of other generators.

**Figure 5.**
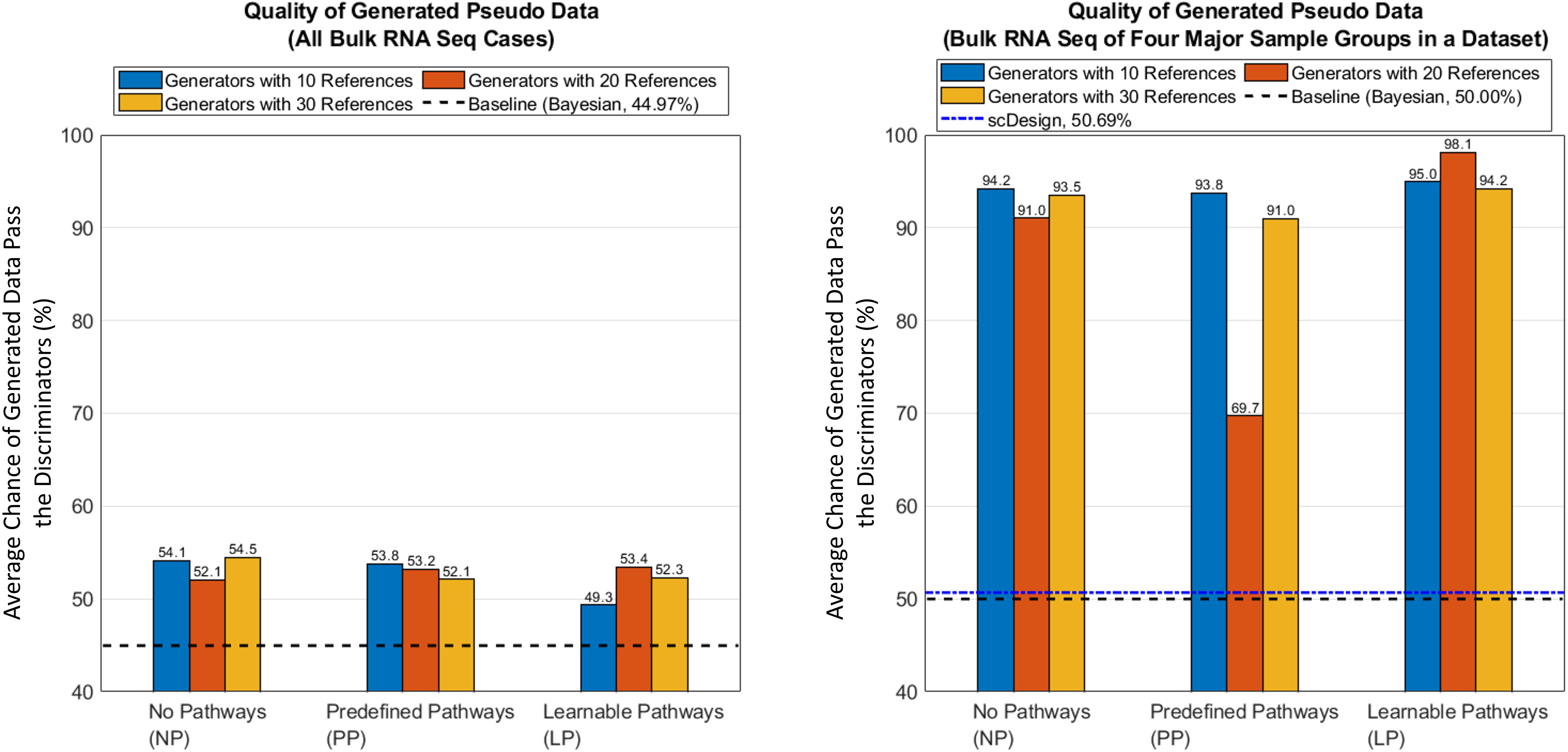
The average chances that pseudo data pass the discrimination test compared to the base line(s) in single-cell and bulk sequencing samples. The values for paired discriminator against generator were not suitable for comparison because they have interacted with each other during the training procedure, so the corresponding values have been replaced with the averaged score of other generators.

### Function 4: Vectorization

The generation of a vector data set is available by using Generators. The provision of a set of targeted samples to the generator and collection of the vector in the latent-space layer enables the acquisition of a 64-dimensional vector corresponding to the targeted sample group, thereby encapsulating the essential features. The dimension is set with consideration of the amount of protentional cell/sample types and the representation capacity, as it is crucial to ensure the quality of pseudo data [49]. This vector can be utilized for a variety of purposes, including sample clustering and visualization, similarity search, and transfer learning acceleration of models based on RNAGAN.

The ideal vectorization of samples can be characterized by two fundamental properties: the maintenance of major informative features, and the elimination of irrelevant variances or biological noise. To validate the performance of RNAGAN in this aspect, samples were generated for the bulk-sequencing cases with more than 30 samples. The latent-space vectors of the NP30 generator (which preformed the best against all discriminator versions) were then compared with those of other commonly adopted clustering and visualization approaches (PCA, T-SNE, and UMAP) (**Figure 6**).

**Figure 6.**
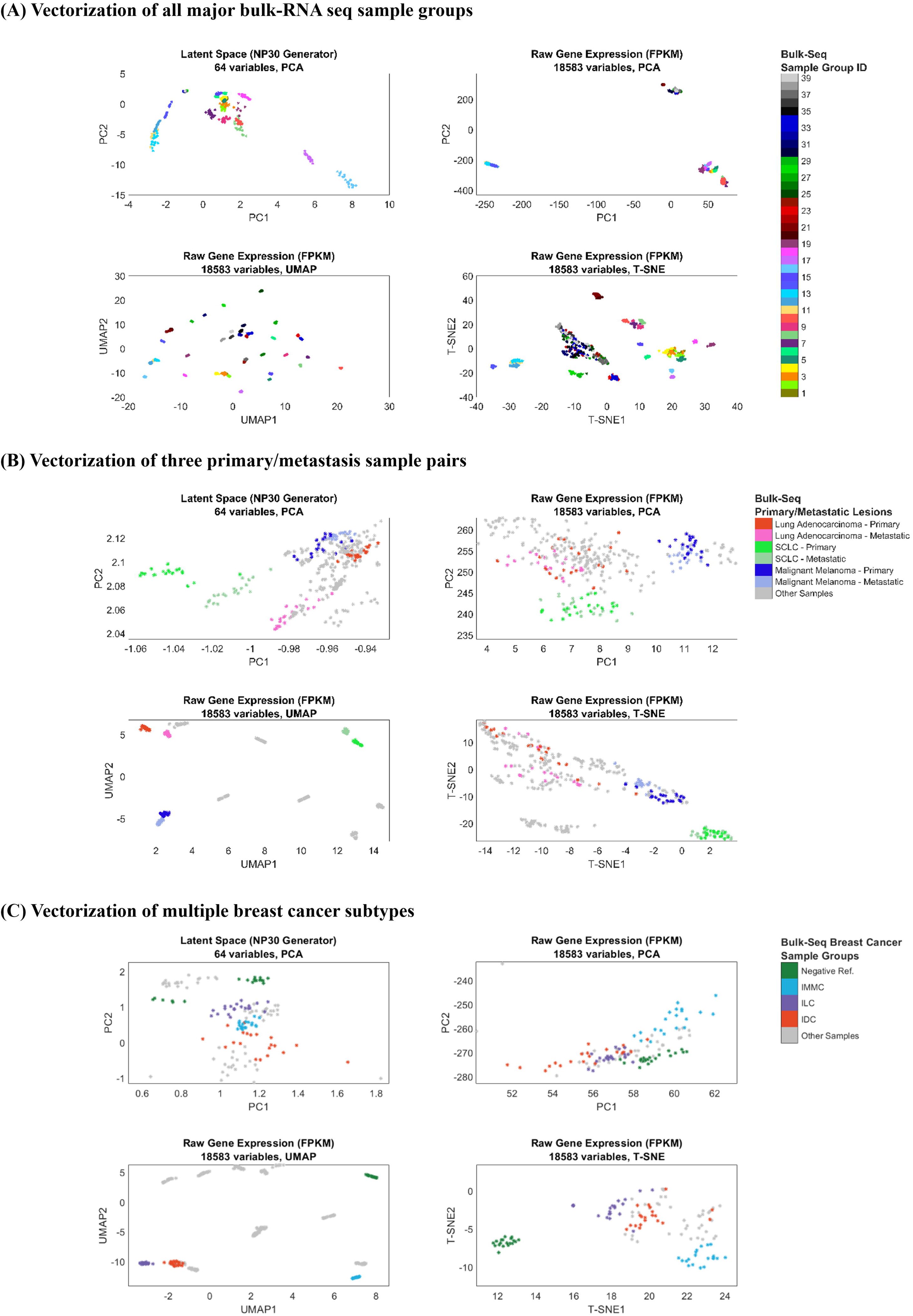
Vectorization of bulk-RNA sequencing sampling groups with over 30 available samples. (**A**) results of all groups. (**B**) results of primary/metastasis sample pairs. (**C**) multiple breast cancer subtypes.

It is evident that the direct PCA of all the genes and T-SNE are inadequate in accurately distinguishing all the sample groups (**Figure 6A**). In the vectorization of paired primary and metastatic lesions of three cancer types (**Figure 6B**), both the latent space of RNAGAN and UMAP have been demonstrated to effectively separate paired samples and accurately locate them based on their similarity. In contrast, direct PCT and T-SNE methods have been found to lack the capacity to clearly differentiate between lesions. Although UMAP can successfully differentiate the samples, the distance from UMAP may not accurately reflect the intragroup variations. As in the test of vectorizing multiple breast cancer subtypes (**Figure 6C**), there are multiple irrelevant sample groups that appeared between the breast cancer groups in UMAP. Conversely, no similar problems have been found in RNAGAN results. In summary, RNAGAN vectors offer the advantages of traditional dimension reduction methods in terms of grouping and similarity-distance relationships, while demonstrating enhanced robustness in their performance.

## Discussions

We presented RNAGAN, a generative adversarial network structure to process single-cell and bulk RNA sequencing, for the following four purposes.

1. Establish cell/sample level stratification or diagnosis with 20-30 positive reference samples;
2. Interpretation of the stratification at the gene / path level, identify related marker genes and the protentional regulating mechanisms among them;
3. Generate high quality pseudo samples for data enrichment or sample representativeness, with structurally preventing memorization of individual real samples;
4. Search for similar sample types with the corresponding vector database, which can be used to track cell lineage or identify similar samples that share similar mechanisms.

The practice of this study aligns with multiple recent recommendations in the field of AI: the avoid of feature leakage between known different objects (such as human vs. mouse [21]), continuously improving the performance of AI by utilizing the completions between subnetworks [50], and multiple suggested “tricks” in network design and training [29, 32].

In the validation of the diagnosis of the sample, RNAGAN achieved AUC above 80% in the diagnosis of multiple common cancer types. This advantage arises from the pathway-embedded design and pseudo-sample generation, which enhance representativeness even with 20 to 30 available reference samples, which is suitable for clinical application. This can also be a reference for future zero-short diagnosis, which would be a significantly more challenging task to achieve accurate diagnosis with less than 20 positive samples compared to more than 30 samples.

RNAGAN has a unique network layer to extract pathway activities, which thus lead to three major network versions: with predefined pathways (PP version) from the Human Molecular Signatures Database, MSigDB, v2023.1.Hs [26, 27, 30], with learned pathways (LP version), and without pathway activity layer (NP version). The pathway layer and predefined pathways are beneficial in the diagnosis of many types of cancer. However, in contrast to Alpha-zero, the LP version does not demonstrate a substantial advantage in terms of outcompeting the other two versions. For scenarios where LP offers adequate diagnosis accuracies, either NP or PP (or both) could also attain comparable or higher levels of accuracy. Consequently, the LP version does not demonstrate exceptional performance in this study.

RNAGAN can extract meaningful features to support or explain its results, which can help researchers better understand the results and provide biologically meaningful insights into the disease. We extracted marker genes and pathways of 13 types of cancer (types with more than 30 positive samples in the datasets) and identified makers and mechanisms that consists with current studies on these diseases. And it should be emphasized that these results are purely data-driven, and further analysis and experiments are compulsory before clinical applications.

Regarding the generation of synthetic data, we compared RNAGAN with other currently adopted approaches, and the generated synthetic data is more “real” in terms of passing the discriminators of RNAGAN.

### Limits and the Next Step

The current version of RNAGAN focused only on the RNA sequencing of 18,583 coding genes. Although current networks already achieved AUC over 80% in the validations of representative cancer types, similar networks can be established for multi-omics data and may obtain more information for the diagnosis and mechanism interpretation of diseases.

Moreover, AI is one of the most active plants, and there are hundreds of new structures or mechanisms every year that are proposed, such as transformer models [51] and nested learning [52]. The adoption of these achievements may also significantly improve the performance of networks and provide more insight for medical researchers.

The project has commenced the subsequent phase of validation, with a focus on external datasets, with a particular emphasis on rare and regional diseases. It is imperative and constitutes the ultimate objective in the development of this network structure.

### Contributions for Medical AI Networks

In this study, we proposed our RNAGAN network, not only for the purpose of its four features, but also to highlight the remarkable presentation of the concept of Generative Adversarial Networks (GAN), which has not yet been fully expanded in previously [53]. Benefited from the internal competitive structure (generator and discriminator), different sub-networks can be easily adopted for multiple different medical challenges while maintaining their highly targeted functionality, which is very different from common single-task agents and generally purpose LLM approaches, and could structurally provide better interoperability for validation and insight generation.

The proposed network also showed a practical approach to structurally embed current biomedical knowledge into an AI network and lead to significant benefits. Such as the Model-Based Transfer Learning (MBTL) procedure that achieves models for both single-cell and bulk level applications. The anonymization layers also show a practice that structurally prevents the leakage of data of certain samples. These designs and concepts could be adopted in other AI networks.

For the purpose of cell/sample stratification, this study shows that 20-30 positive referencing samples could be a sufficient sample size for most of the testing cancer types. Although future analysis tools may reduce the sample size with comparable accuracies, they would require reasonably deep biomedical insights.

## Author contributions

WD supervised this study. ZH developed the study methodology and contributed to data acquisition and analysis. ZH contributed to data interpretation. WD and ZH acquired the funding for this project. All authors contributed to writing, reviewing, and revising the manuscript.

## Acknowledgements

This work was supported by the Hong Kong Health and Medical Research Fund (11221746) (to ZH) and Theme-based Research Scheme (T12-703/22-R; T123-70323-N) from the Research Grant Council (to WD).

## Data Sharing Statement

The data for network training and validation comes from multiple public databases, which have been summarized in supplementary **Table S1** and **S2**. RNAGAN, the trained network and relative software scripts are available on https://github.com/ZhaozhengHou-HKU/RNAGAN-1.0.git.

## Supplementary figures and tables

**Figure S1**. Extracted Features of breast cancer ILC and classical GBM.

**Table S1**. Details of used single-cell RNA sequencing datasets.

**Table S2**. Details of used bulk RNA sequencing datasets.

**Table S3**. Evaluation of the diagnosis of 13 major cancer types.

**Table S4**. Extracted maker genes/pathways of the major cancer types.

**Table S5**. List of predefined pathways and the mapping of learned pathways from LP discriminator networks to known gene pathways with identified biomedical meanings.

## Notes

https://github.com/ZhaozhengHou-HKU/RNAGAN-1.0.git

### Competing Interest Statement

The authors have declared no competing interest.

