## Supplementary figures and images for "RNAGAN: Train One and Get Four, Multipurpose Human RNA-Seq Analysis Tool with Enhanced Interpretability and Small Data Size Capability"

### Figure S1

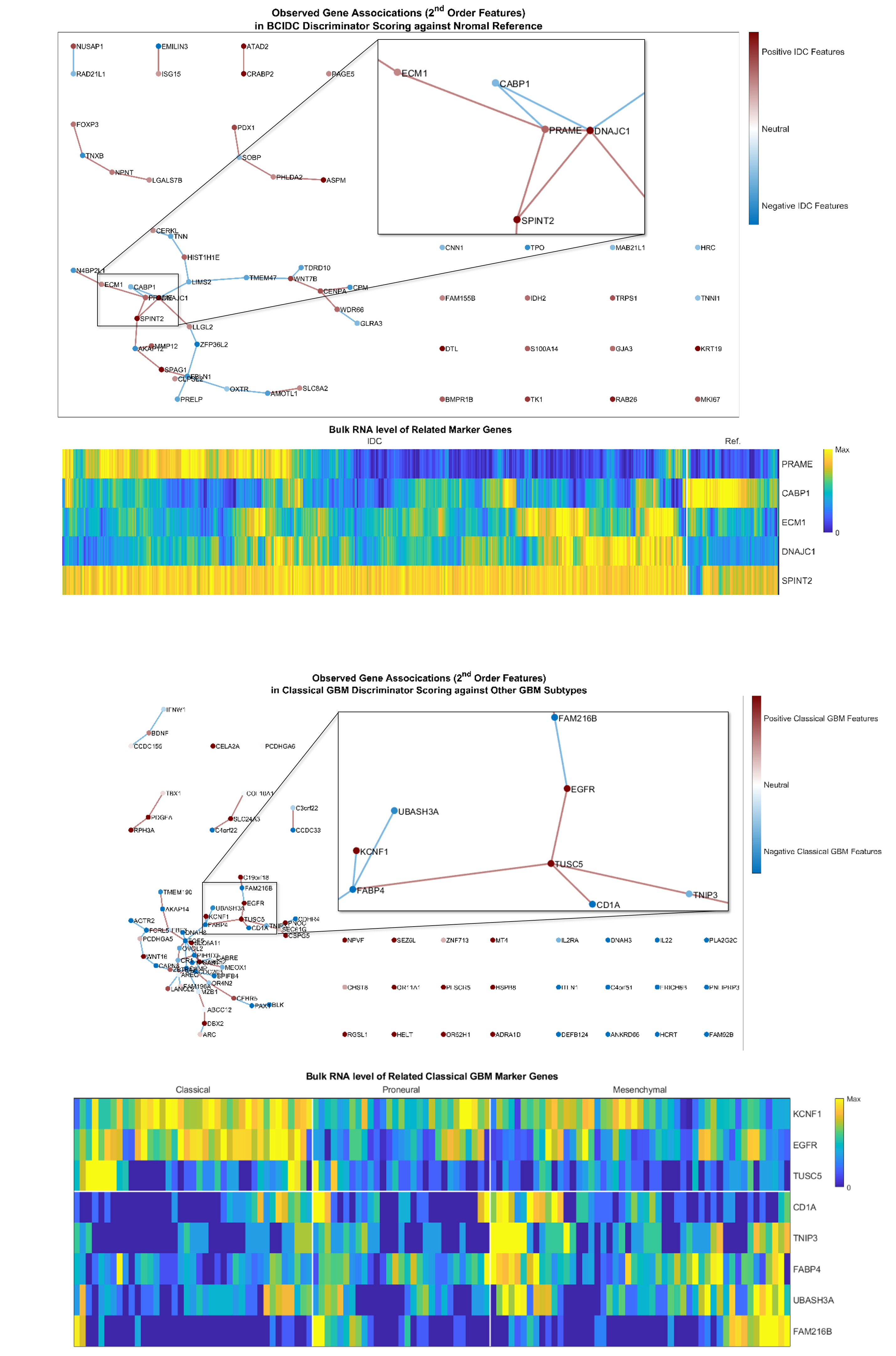
