## Supplementary material for "RNAGAN: Train One and Get Four, Multipurpose Human RNA-Seq Analysis Tool with Enhanced Interpretability and Small Data Size Capability": Table S1

Non-commercial preprint on bioRxiv.

**The authors declare no potential conflicts of interest.**

### Supplementary tables

Table S1. Single cell datasets

| Index | Publication (Authors) | Year | # of cells | Assay | Diseases or Tissue |
|---|---|---|---|---|---|
| **1** | Nowicki-Osuch, et al. [1] | 2023 | 146,583 | 10x 3' v2<br>10x 3' v3 | barrett esophagus, gastric intestinal metaplasia, gastritis |
| **2** | Salcher, et al. [2] | 2022 | 892,296 | 9 assays | chronic obstructive ulmonary disease, lung adenocarcinoma, non-small cell lung carcinoma, squamous cell lung carcinoma |
| **3** | Han, et al. [3] | 2020 | 599,926 | microwell-seq | normal multitissue |
| **4** | Lake, et al. [4] | 2023 | 107,344 | 10x 3' v3 | acute kidney failure, chronic kidney disease |
| **5** | Cerami, et al. [5] | 2012 | 186,887 | 10x 3' v2 | esophageal squamous cell carcinoma, esophageal adenocarcinoma |
| **6** | Domínguez Conde, et al. [6] | 2022 | 329,762 | 10x 3' v3<br>10x 5' v1<br>10x 5' v2 | normal multitissue |
| **7** | Kumar, et al. [7] | 2023 | 650,986 | 10x 3' v2<br>10x 3' v3 | normal breast |
| **8** | Eraslan, et al. [8] | 2022 | 209,126 | 10x 3' v2 | normal multitissue |
| **9** | Bhat-Nakshatri, et al. [9] | 2021 | 31,696 | 10x 3' v2<br>10x 3' v3 | normal breast |
| **10** | Andrews, et al. [10] | 2022 | 68,792 | 10x 3' v2<br>10x 3' v3 | normal liver |
| **11** | Sikkema, et al. [11] | 2023 | 584,944 | 5 assays | normal lung |
| **12** | Consortium*, et al. [12] | 2022 | 416,602 | 10x 3' v3<br>10x 5' v2<br>Smart-seq2<br>Smart-seq3 | normal multitissue |
| **13** | Fasolino, et al. [13] | 2022 | 69,645 | 10x 3' v2<br>10x 3' v3 | diabetes mellitus |
| **14** | Ruiz-Moreno, et al. [14] | 2022 | 338,564 | 8 assays | glioblastoma |

Table S2. Bulk sequencing datasets

| Index | Publication (Authors) | Year | # of samples | Diseases or Tissue |
|---|---|---|---|---|
| **1** | Ciriello, et al. [15] | 2015 | 1,084 | breast invasive carcinoma (nos/ ductal carcinoma/ lobular carcinoma/ mixed mucinous carcinoma) |
| **2** | Brennan, et al. [16] | 2013 | 577 | glioblastoma (classical/ g-cimp/ mesenchymal/ neural/ proneural) |
| **3** | Bonneville, et al. [17] | 2017 | 594 | colon adenocarcinoma, rectal adenocarcinoma, mucinous adenocarcinoma of the colon and rectum |
| **4** | Network [18] | 2014 | 566 | non-small cell lung cancer (lung adenocarcinoma) |
| **5** | Liu, et al. [19] | 2018 | 372 | hepatocellular carcinoma |
| **6** | Liu, et al. [19] | 2018 | 523 | head and neck squamous cell carcinoma |
| **7** | Bailey, et al. [20] | 2016 | 456 | pancreatic cancer |
| **8** | Ghandi, et al. [21], Nusinow, et al. [22] | 2019, 2020 | 1,739 | carcinoid-endocrine tumour, chondrosarcoma, fibrosarcoma, giant cell tumour, glioma, haematopoietic neoplasm, leiomyosarcoma, lymphoid neoplasm, malignant melanoma, meningioma, mesothelioma, neuroblastoma, osteosarcoma, rhabdoid tumour, rhabdomyosarcoma, sex cord-stromal tumour, and others |

Table S3. Evaluation of the diagnosis of 13 major cancer types.

| AUC | | No Pathways (NP) | | | Predefined Pathways (PP) | | | Learnable Pathways (LP) | | |
|---|---|---|---|---|---|---|---|---|---|---|
| | | 10 Ref | 20 Ref | 30 Ref | 10 Ref | 20 Ref | 30 Ref | 10 Ref | 20 Ref | 30 Ref |
| overall AUC | | 77.53% | 82.69% | 78.39% | 81.29% | 79.53% | 82.34% | 84.98% | 83.24% | 82.09% |
| Target Cancer Type and Number of references | Negative Samples | 73.5% - 80.7% | 79.3% - 85.5% | 75.1% - 81.3% | 78.7% - 84.2% | 76.4% - 83.1% | 78.8% - 85.0% | 58.0% - 87.6% | 79.5% - 85.8% | 78.3% - 84.3% |
| IDC, Breast Invasive Ductal Carcinoma | Negative Reference | 51.68% | 92.83% | 19.39% | 97.56% | 95.74% | 97.87% | 51.53% | 26.21% | 16.20% |
| 232 | 114 | 44.5% - 57.6% | 88.3% - 96.2% | 13.9% - 27.2% | 94.1% - 98.9% | 92.2% - 97.6% | 95.5% - 99.3% | 43.0% - 60.3% | 19.6% - 32.7% | 11.8% - 20.2% |
| ILC, Breast Invasive Lobular Carcinoma | Negative Reference | 54.78% | 94.51% | 20.41% | 92.68% | 96.22% | 92.42% | 38.02% | 32.20% | 10.71% |
| 60 | 114 | 46.8% - 61.9% | 88.2% - 97.3% | 14.5% - 26.5% | 86.1% - 95.3% | 92.0% - 97.8% | 86.5% - 95.6% | 31.3% - 47.2% | 25.1% - 40.3% | 6.9% - 16.3% |
| Classical GBM | Other GBM subtypes | 69.36% | 62.85% | 77.30% | 53.99% | 40.94% | 53.03% | 42.04% | 70.53% | 66.18% |
| 11 | 104 | 61.7% - 76.8% | 56.1% - 73.3% | 69.1% - 83.2% | 44.1% - 60.3% | 34.0% - 47.7% | 42.1% - 57.8% | 35.3% - 50.6% | 63.3% - 77.9% | 57.7% - 74.0% |
| Mesenchymal GBM | Other GBM subtypes | 37.90% | 37.39% | 35.90% | 53.72% | 56.27% | 48.63% | 55.99% | 36.66% | 50.46% |
| 14 | 94 | 31.3% - 48.1% | 29.8% - 49.6% | 28.8% - 42.3% | 45.2% - 62.2% | 45.5% - 65.9% | 40.5% - 54.9% | 48.4% - 64.7% | 30.6% - 43.3% | 43.1% - 56.2% |
| Neural GBM | Other GBM subtypes | 34.10% | 47.53% | 34.16% | 35.37% | 70.38% | 66.66% | 62.48% | 45.54% | 53.48% |
| 9 | 117 | 25.6% - 43.1% | 40.2% - 53.9% | 25.9% - 41.4% | 29.1% - 41.8% | 64.5% - 77.6% | 57.5% - 73.7% | 55.7% - 72.6% | 35.9% - 54.7% | 45.8% - 62.1% |
| Proneural GBM | Other GBM subtypes | 48.93% | 73.89% | 51.56% | 39.76% | 50.20% | 54.98% | 50.98% | 52.30% | 53.05% |
| 9 | 114 | 41.3% - 56.3% | 64.8% - 80.1% | 42.6% - 60.5% | 30.6% - 47.0% | 42.8% - 58.7% | 48.2% - 63.7% | 44.2% - 57.1% | 45.9% - 63.7% | 46.7% - 61.1% |
| COAD, Colon Adenocarcinoma | Negative Reference | 24.80% | 95.63% | 3.54% | 99.49% | 95.53% | 97.96% | 57.76% | 93.58% | 99.43% |
| 113 | 51 | 18.1% - 34.5% | 91.1% - 97.7% | 1.5% - 6.1% | 98.1% - 99.9% | 92.3% - 98.1% | 94.2% - 99.3% | 49.2% - 66.2% | 89.8% - 96.6% | 97.8% - 100.0% |
| READ, Rectal Adenocarcinoma | Negative Reference | 23.22% | 97.01% | 0.53% | 100.00% | 98.26% | 99.87% | 67.34% | 97.46% | 100.00% |
| 40 | 51 | 17.1% - 31.0% | 93.5% - 99.1% | 0.2% - 1.4% | 100.0% - 100.0% | 96.5% - 99.3% | 99.4% - 100.0% | 59.9% - 72.4% | 93.6% - 99.4% | 100.0% - 100.0% |
| HCC, Hepatocellular Carcinoma | Negative Reference | 49.59% | 21.03% | 11.65% | 99.26% | 94.36% | 99.12% | 76.39% | 86.90% | 96.17% |
| 106 | 50 | 40.3% - 55.8% | 16.8% - 26.5% | 8.2% - 16.2% | 97.9% - 99.9% | 87.0% - 96.8% | 97.5% - 99.8% | 68.3% - 82.1% | 80.2% - 90.6% | 93.2% - 97.7% |
| HNSC, Head and Neck Squamous Cell Carcinoma | Negative Reference | 15.53% | 76.98% | 19.16% | 41.45% | 96.36% | 74.70% | 75.34% | 33.23% | 87.92% |
| 151 | 44 | 11.1% - 22.2% | 67.2% - 83.0% | 14.5% - 25.2% | 30.7% - 47.9% | 93.4% - 98.0% | 68.3% - 81.6% | 67.7% - 81.8% | 25.6% - 41.5% | 81.7% - 92.5% |
| Acute Myeloid Leukaemia | Other cancer types | 92.76% | 96.80% | 82.28% | 19.05% | 56.13% | 30.96% | 32.72% | 42.18% | 25.45% |
| 9 | 407 | 87.3% - 95.8% | 94.2% - 98.4% | 75.9% - 88.4% | 13.8% - 26.4% | 47.6% - 65.1% | 24.2% - 37.8% | 27.6% - 41.0% | 35.9% - 52.4% | 19.5% - 35.1% |
| Large Intestine Adenocarcinoma | Other cancer types | 51.18% | 19.89% | 29.39% | 51.01% | 58.86% | 52.27% | 52.75% | 38.16% | 42.80% |
| 10 | 404 | 40.7% - 57.1% | 14.4% - 25.8% | 23.2% - 36.1% | 41.4% - 58.1% | 50.7% - 69.3% | 44.0% - 60.7% | 42.9% - 59.2% | 28.9% - 46.1% | 35.8% - 51.5% |
| Malignant Melanoma | Other cancer types | 49.17% | 72.17% | 76.83% | 62.50% | 73.91% | 63.50% | 74.23% | 74.46% | 47.56% |
| 10 | 407 | 40.0% - 58.2% | 63.5% - 79.3% | 69.1% - 83.8% | 56.6% - 69.9% | 65.3% - 80.5% | 56.2% - 70.6% | 67.3% - 79.4% | 68.1% - 80.9% | 39.6% - 56.1% |
